# Distinct Mechanisms of Type 3 Secretion System Recognition Control LTB_4_ Synthesis in Neutrophils versus Macrophages

**DOI:** 10.1101/2024.07.01.601466

**Authors:** Amanda Brady, Leonardo C. Mora-Martinez, Benjamin Hammond, Bodduluri Haribabu, Silvia M. Uriarte, Matthew B. Lawrenz

## Abstract

Leukotriene B4 (LTB_4_) is critical for initiating the inflammatory cascade in response to infection. However, *Yersinia pestis* colonizes the host by inhibiting the timely synthesis of LTB_4_ and inflammation. Here, we show that the bacterial type 3 secretion system (T3SS) is the primary pathogen associated molecular pattern (PAMP) responsible for LTB_4_ production by leukocytes in response to *Yersinia* and *Salmonella*, but synthesis is inhibited by the Yop effectors during *Yersinia* interactions. Moreover, we unexpectedly discovered that T3SS-mediated LTB_4_ synthesis by neutrophils and macrophages require two distinct host signaling pathways. We show that the SKAP2/PLC signaling pathway is essential for LTB_4_ production by neutrophils but not macrophages. Instead, phagocytosis and the NLRP3/CASP1 inflammasome are needed for LTB_4_ synthesis by macrophages. Finally, while recognition of the T3SS is required for LTB_4_ production, we also discovered a second unrelated PAMP-mediated signal independently activates the MAP kinase pathway needed for LTB_4_ synthesis. Together, these data demonstrate significant differences in the signaling pathways required by macrophages and neutrophils to quickly respond to bacterial infections.

**Significance:** The production of inflammatory lipid mediators by the host is essential for timely inflammation in response to invasion by bacterial pathogens. Therefore, defining how immune cells recognize pathogens and rapidly produce these lipids is essential for us to understand how our immune system effectively controls infection. In this study, we discovered that the host signaling pathways required for leukotriene B4 (LTB_4_) synthesis differ between neutrophils and macrophages, highlighting important differences in how immune cells respond to infection. Together, these data represent a significant improvement in our understanding of how neutrophils and macrophages rapidly react to bacteria and provide new insights into how *Yersinia pestis* manipulates leukocytes to evade immune recognition to cause disease.

## Introduction

Rapid recruitment of immune cells to sites of infection is essential to control bacterial pathogens. However, sustained or chronic inflammation is detrimental to the host. Therefore, inflammation is a tightly regulated cascade of events controlled by both lipid and protein mediators (1). Initiation of this cascade is primarily mediated by inflammatory lipids, of which the eicosanoid leukotriene B4 (LTB_4_) is one of the earliest produced during infection. LTB_4_ is a potent pro-inflammatory chemoattractant and immune cell activator necessary for timely recruitment of neutrophils to sites of infections (1–6). LTB_4_ is derived from arachidonic acid (AA) upon activation of the enzymes cytosolic phospholipase A_2_ (cPLA_2_) and 5-lipoxygenase (5-LOX). The enzymes are activated by phosphorylation via MAPK signaling and Ca^2+^ binding to translocate to the nuclear membrane or a lipidisome, where a complex is formed with 5-LOX activating protein (FLAP) (7–14). This complex converts AA to LTA_4_, which is rapidly converted to LTB_4_ by LTA_4_ hydrolase (7, 8). Upon subsequent release, LTB_4_ is recognized by the high affinity BLT1 receptor on immune cells to promote priming, activation, and chemotaxis. Mouse models that disrupt the host’s ability to produce or respond to LTB_4_ are impaired in initiating inflammation and are significantly more susceptible to a variety of infections, demonstrating the importance of LTB_4_ in inducing a robust inflammatory response required for effective clearance of pathogens (1, 2, 15, 16).

Plague is an acute infection caused by the bacterial pathogen *Yersinia pestis*. A hallmark manifestation of plague is a significant delay in the recruitment of immune cells to the sites of infection, allowing for bacterial colonization and replication (17–20). One of the key virulence determinants for *Y. pestis* to inhibit the immune system and colonize the host is the Ysc type 3 secretion system (T3SS) encoded on the pCD1 plasmid (21, 22). This secretion system mediates direct translocation of bacterial effector proteins called Yops into host cells (22–25). During mammalian infection, *Y. pestis* primarily targets neutrophils and macrophages for T3SS-mediated injection of the Yops (26–28). Once inside host cells, the Yop effectors target specific host factors to inhibit phagocytosis, reactive oxygen species (ROS) synthesis, degranulation by neutrophils, and inflammatory cytokine and chemokine release that is required to recruit circulating leukocytes to infection sites (29–37). Moreover, the Yop proteins were recently shown to actively inhibit the synthesis of LTB_4_ required for the initiation of the inflammatory cascade during pneumonic plague (37, 38). However, in the absence of the Yop effectors, neutrophils interacting with *Y. pestis* can synthesize LTB_4_, but synthesis requires bacterial expression of the T3SS and the YopB/D translocase (38). These data indicate that the T3SS is a pathogen associated molecular pattern (PAMP) recognized by neutrophils that triggers signaling pathways that lead to inflammatory lipid synthesis.

Previous studies in macrophages have established components of the T3SS induce inflammatory cell death pathways. In the absence of the Yops, the T3SS and YopB/D translocon proteins can induce NLRP3-dependent activation of the caspase 1 inflammasome, IL1-β secretion, and pyroptosis, which is inhibited by YopK (31, 39). These studies suggest that T3SS-dependent LTB_4_ synthesis by neutrophils may also require NLRP3/CASP1 inflammasome activation. However, LTB_4_ synthesis in response to other stimuli is not dependent on inflammasome activation (11, 40, 41). Furthermore, infection of neutrophils with a strain of *Y. pestis* that lacks all the Yops except YopK does not inhibit LTB_4_ synthesis in neutrophils (37, 38), raising an alternative possibility for inflammasome-independent mechanisms leading to T3SS-dependent LTB_4_ synthesis. Moreover, Yop effectors other than YopK that have not been directly linked to inflammasome inhibition – YpkA, YopE, YopJ, or YopH – are sufficient to independently inhibit LTB_4_ synthesis by neutrophils (37, 38), further supporting the latter hypothesis. Of these four Yop effectors, YopJ and YopH are intimately involved in MAPK and Ca^2+^ signaling required for LTB_4_ synthesis. YopJ is an acyltransferase that targets several kinases in the MAPK pathway (42–45), and Pulsifer et al. demonstrated that YopJ inhibition of ERK phosphorylation is sufficient to inhibit LTB_4_ synthesis by human neutrophils (37). YopH is a tyrosine phosphatase that has been shown to target multiple proteins of the focal adhesion complex, including SKAP2, SLP-76, PRAM, Vav, and LCK (28, 46–49). Through these interactions, YopH inhibits β1-integrin-mediated Ca^2+^ flux by neutrophils during interactions with *Y. pseudotuberculosis*, raising the possibility that this kinase signaling hub might be involved in LTB_4_ synthesis in response to the T3SS. In this study, we use *Y. pestis* Yop mutants to define the molecular mechanisms responsible for T3SS-dependent LTB_4_ synthesis and discovered significant differences in the host factors required for T3SS-sensing and LTB_4_ synthesis between neutrophils and macrophages.

## Results

### Neutrophil LTB_4_ synthesis in response to *Salmonella enterica* Typhimurium is dependent on the SPI-1 T3SS

LTB_4_ synthesis in response to *Y. pestis* is dependent on the expression of the T3SS and the YopB/D translocon (38). Moreover, *S. enterica* Typhimurium, which encodes two T3SSs (SPI-1 and SPI-2), also induces an LTB_4_ response in neutrophils, but whether synthesis required expression of the T3SSs was not tested (50). To determine if leukocyte sensing of the T3SSs of *S. enterica* Typhimurium was responsible for LTB_4_ synthesis, bone marrow derived murine neutrophils (BMNs) were infected with *S. enterica* Typhimurium LT2 (ST+) or a mutant lacking both the *Salmonella* pathogenicity island 1 (SPI-1) and SPI-2 encoded type 3 export apparatuses (ΔSPI1/2). After 1 h of infection, LTB_4_ synthesis was significantly elevated in ST+ infected BMNs (Fig 1A, ST+ vs. UI, p≤0.0001) but it was not elevated in cells infected with the ΔSPI1/2 mutant (Fig 1A, p= 0.1513). To determine the contribution of the individual T3SSs, BMNs were next infected with individual SPI-1 or SPI-2 mutants. No significant differences in recovered LTB_4_ were observed between cells infected with the ΔSPI1/2 and ΔSPI1 mutant strains, but LTB_4_ concentrations were significantly elevated in the ΔSPI2 mutant infected cells (Fig 1A, ΔSPI1/2 vs. ΔSPI2, p≤0.0001). Together these data support that neutrophils not only sense the *Y. pestis* T3SS but also the SPI-1 T3SS of *S. enterica* Typhimurium to initiate a robust LTB_4_ response.

**Fig. 1.**
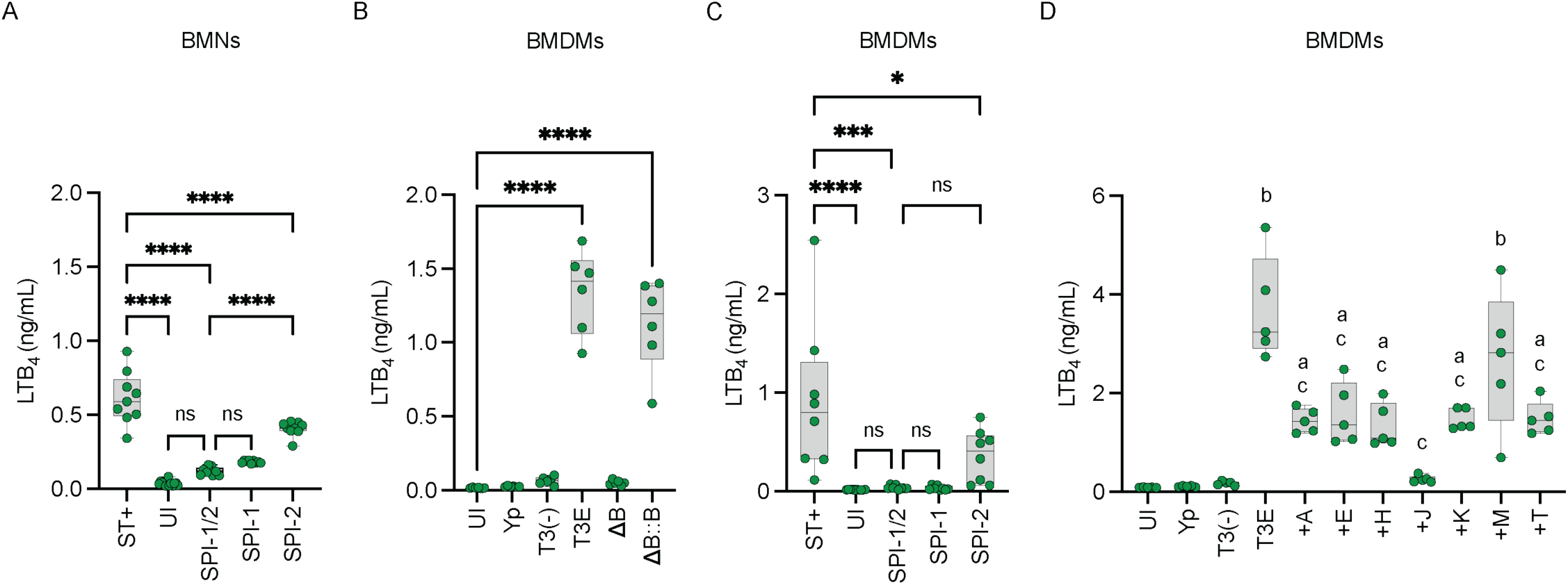
T3SS-depedent LTB_4_ synthesis is conserved in *S. enterica* Typhimurium and BMDMs. (A) Murine neutrophils (BMNs) or (C) Murine macrophages (BMDMs) were infected with *S*. *enterica* Typhimurium LT2 (ST+), mutants that lacked *Salmonella* T3SS SPI-1, SPI-2, or both SPI-1/2. (B) BMDMs were infected with *Y. pestis, Y. pestis* T3E, *Y. pestis* T3^(−)^, a yopB mutant in the T3E background (ΔB), or ΔB complemented with yopB (ΔB::B). (D) BMDMs infected with *Y. pestis, Y. pestis* T3E, *Y. pestis* T3^(−)^, or *Y. pestis* strains expressing only one Yop (+A = YpkA; +E = YopE; +H = YopH; +J = YopJ; +K = YopK; +M = YopM; or +T = YopT). (A) BMNs were infected at an MOI of 20 for 1 h. (B-D) BMDMs were infected at an MOI of 20 for 4 h. (A-D) LTB_4_ was measured from supernatants by ELISA. Each symbol represents an independent biological infection, and the box plot represents the median of the group ± the range. UI = uninfected. ns = not significant. One-way ANOVA with Tukey’s *post hoc* test compared to each condition for A, C, and D, or Dunnett’s *post hoc* test compared to uninfected for B. *=p≤0.05, ***=p≤0.001, ****=p≤0.0001. For panel D, p values when compared to uninfected denoted as a=p≤0.05 or b=p≤0.0001, and when compared to *Y. pestis* T3E as c=p≤0.0001.

### Requirement for interactions with the YopB/D translocase for LTB_4_ synthesis is conserved in macrophages

While we have previously shown that LTB_4_ synthesis by both neutrophils and M1-polarized macrophages in response to *Y. pestis* is dependent on the presence of the T3SS, we only demonstrated a requirement for the YopB/D translocase in neutrophils (38). To determine whether the YopB/D translocase is also required for LTB_4_ synthesis by macrophages, M1-polarized bone marrow derived macrophages (BMDMs) were infected with *Y. pestis*, a *Y. pestis* strain expressing the T3SS but lacking all Yop effectors (*Y. pestis* T3E), a *Y. pestis* strain lacking the pCD1 plasmid encoding the entire Ysc T3SS [*Y. pestis* T3^(−)^], or a *Y. pestis* T3E *yopB* mutant that is defective in expression of the translocase that directly interacts with the host cell plasmid membrane (51–53). As previously observed for neutrophils (38), BMDMs also did not synthesize LTB_4_ in response to *Y. pestis* T3^(−)^ or *Y. pestis* T3E *yopB* (Fig 1B). Normal LTB_4_ synthesis was restored by *yopB* complementation (Fig 1B, *yopB*::c*yopB*). To confirm that the *S. enterica* Typhimurium SPI-1 T3SS is also required for LTB_4_ synthesis by macrophages, BMDMs were infected with ST+ or the ΔSPI1/2, ΔSPI1, or ΔSPI2 mutants. As observed for BMNs, BMDM synthesis of LTB_4_ was dependent on the presence of the SPI-1 T3SS but not SPI-2 system (Fig 1C). These data show that like neutrophils, macrophages respond to bacterial T3SSs by rapidly synthesizing LTB_4_.

### Only YopJ is sufficient to inhibit LTB_4_ synthesis by macrophages

We have previously shown that YpkA, YopE, YopJ, or YopH are individually sufficient to inhibit LTB_4_ synthesis in neutrophils (37, 38). To determine if these Yop effectors could inhibit LTB_4_ synthesis by macrophages, LTB_4_ was measured from BMDMs infected with *Y. pestis* strains that expressed only one Yop effector. BMDMs infected with strains expressing YpkA, YopE, YopH, YopK, and YopT showed significant decreases in LTB_4_ compared to those infected with *Y. pestis* T3E, but still produced significantly more LTB_4_ than *Y. pestis* infected or uninfected cells (Fig 1D). However, BMDMs infected with a strain expressing only YopJ appeared to be completely inhibited in their ability to synthesize LTB_4_, producing concentrations similar to those recovered from cells infected with *Y. pestis* expressing all of the Yop effectors (Fig 1D; +J vs. Yp). YopM was the only effector that did not appear to have any impact on LTB_4_ synthesis. When compared to the ability of individual Yop effectors to inhibit LTB_4_ synthesis in neutrophils, these data suggest that the signaling pathways leading to LTB_4_ in response to the T3SS may differ between macrophages and neutrophils.

### Bacterial phagocytosis enhances LTB_4_ synthesis by macrophages but not neutrophils

One important consequence of Yop intoxication of leukocytes is the inhibition of phagocytosis via the action of YpkA and YopE (54, 55). Moreover, an important function of the SPI-1 T3SS of *S. enterica* Typhimurium is to induce phagocytosis (56). Because phagocytosis is required for LTB_4_ synthesis by leukocytes interacting with crystalline silica (11) and *Y. pestis* and the ΔSPI-1 mutant do not elicit LTB_4_ synthesis in both neutrophils and macrophages, we next asked if phagocytosis of *Y. pestis* T3E or ST+ was required for T3SS-dependent LTB_4_ synthesis by treating BMNs and BMDMs with the phagocytosis inhibitor cytochalasin D (cytoD) prior to infection. In BMNs, while treatment with cytoD inhibited phagocytosis of *Y. pestis* T3E (S1 Fig A-B), it did not alter the ability to synthesize LTB_4_ (Fig 2A and B), indicating that Yop-mediated inhibition of phagocytosis is not responsible for the inhibition of T3SS-mediated LTB_4_ synthesis during *Y. pestis* infection in neutrophils. Additionally, as previously reported by Golenkina et al. (50), cytoD treatment increased LTB_4_ synthesis by BMNs infected with ST+ (Fig 2C; p≤0.05), indicating that induction of phagocytosis by *S. enterica* Typhimurium is not required for T3SS-mediated LTB_4_ synthesis. However, in contrast to neutrophils, when phagocytosis was inhibited in BMDMs (S1 Fig C-D), LTB_4_ synthesis was reduced in response to both *Y. pestis* T3E (Fig 2E; p≤0.05) and ST+ (Fig 2F; p≤0.059). Interestingly, LTB_4_ levels were still higher in the cytoD infected BMDMs than the uninfected BMDMs (Fig. 2D). Together, these data suggest that phagocytosis is not required for T3SS-mediated LTB_4_ synthesis by neutrophils but enhances synthesis by macrophages.

**Fig. 2.**
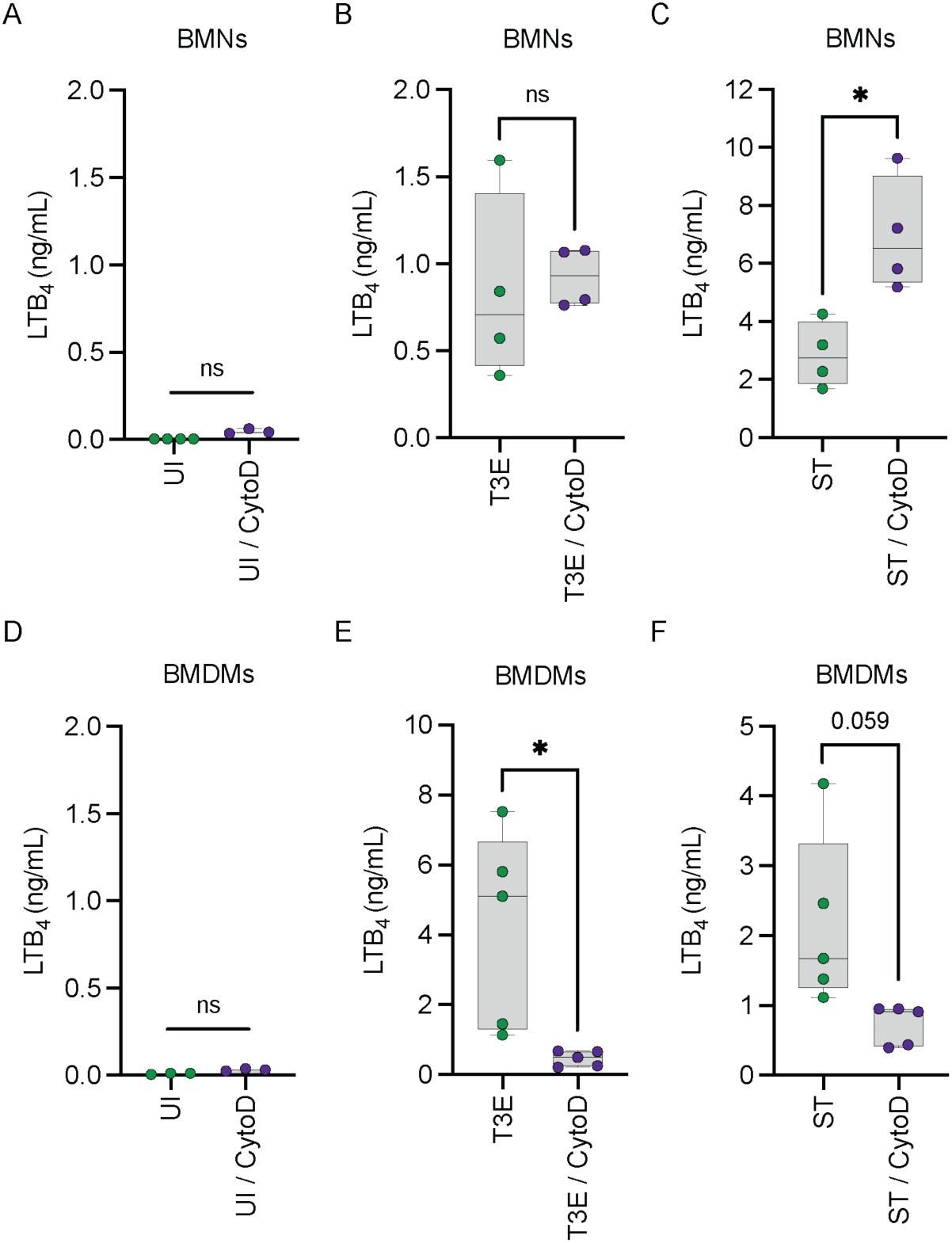
Phagocytosis is not required for LTB_4_ synthesis in BMNs but enhances LTB_4_ synthesis by macrophages in response to *Y. pestis*. (A-C) BMNs or (D-F) BMDMs were untreated (green circles) or pretreated with cytochalasin D (10 μM, purple circles) for 30 min and were either (A,D) uninfected or infected with (B,E) *Y. pestis* T3E or (C,F) ST at an MOI of 20 for (A-C) 1 h or (D-F) 4 h. (A-F) LTB_4_ was measured from supernatants by ELISA. Each symbol represents an independent biological infection, and the box plot represents the median of the group ± the range. UI = uninfected. ns = not significant. T-test with Welch’s *post hoc* test. *=p≤0.05.

### PLC signaling is required for LTB_4_ synthesis in neutrophils but not macrophages

Ca^2+^ flux is required for the activation of cPLA_2_ and 5-LOX (7, 14). However, it is unclear if the T3SS induces Ca^2+^ flux through Ca^2+^ migration through the YopB/D translocase pore or via conventional Ca^2+^ signaling. Phospholipase C (PLC) is the central mediator of conventional Ca^2+^ signaling in the cell (57–60), and chemical inhibitors of PLC have been well characterized (61, 62). Therefore, to determine if Ca^2+^ signaling is required for T3SS-dependent LTB_4_ synthesis, leukocytes were treated prior to infection with U73122, which inhibits PLCβ and PLCγ (61, 63, 64). When PLC signaling was inhibited, BMNs infected with the *Y. pestis* T3E mutant were no longer able to synthesize LTB_4_ compared to untreated BMNs (Fig 3A), suggesting that PLC-mediated Ca^2+^ signaling is required for LTB_4_ synthesis. To ensure that U73122 treatment did not have off target effects on cPLA_2_ or 5-LOX, U73122-treated BMNs were incubated with the Ca^2+^ ionophore, A23187, which induces Ca^2+^ flux and LTB_4_ synthesis independent of PLC signaling (65). Within 10 min of A23187 treatment, U73122-treated BMNs were able to synthesize LTB_4_, supporting that U73122 treatment is not affecting the activity of cPLA_2_ or 5-LOX (Fig 3A). In contrast to BMNs, U73122 treatment of *Y. pestis* T3E-infected BMDMs only modestly inhibited LTB_4_ synthesis compared to untreated cells (Fig 3B; p≤0.05), suggesting PLC is not the primary source of Ca^2+^ flux in macrophages. Together, these data suggest that the T3SS activates PLC-mediated Ca^2+^ flux in neutrophils needed for LTB_4_ synthesis, but alternative mechanisms are required for T3SS-induced LTB_4_ synthesis in macrophages.

**Fig. 3.**
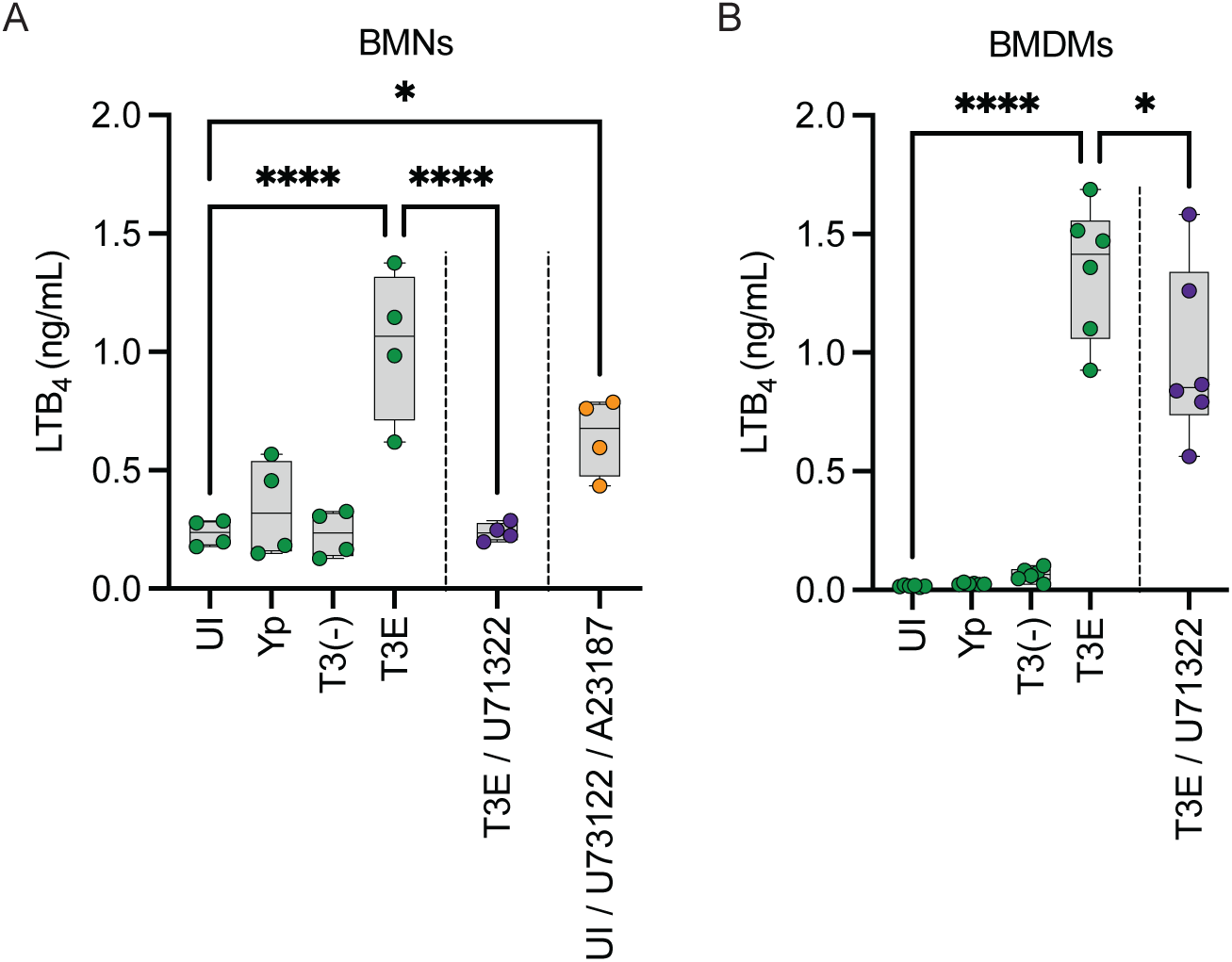
PLC signaling is required for LTB_4_ synthesis in BMNs. (A) BMNs or (B) BMDMs were infected with *Y. pestis, Y. pestis* T3E, or *Y. pestis* T3^(−)^ at an MOI of 20 for (A) 1 h or (B) 4 h. Leukocytes pretreated with PLC inhibitor (U73122; (A) 5 μM or (B) 20 μM) for 30 min (purple circles). (A) BMN supernatants were replaced with fresh media and cells were treated with A23187 (1 μM) for 10 min after treatment with U73122 for 30 min (orange circles). (A-B) LTB_4_ was measured from supernatants by ELISA. Each symbol represents an independent biological infection, and the box plot represents the median of the group ± the range. UI = uninfected. One-way ANOVA with Tukey’s *post hoc* test compared to each condition. *=p≤0.05, ****=p≤0.0001.

### STIM1-mediated flux of extracellular Ca^2+^ is required for LTB_4_ synthesis in both neutrophils and macrophages

PLC-mediated Ca^2+^ flux is required for LTB_4_ synthesis in neutrophils, but PLC signaling can lead to both Ca^2+^ efflux from the ER and influx from the extracellular space (57–60, 66). To determine if intracellular Ca^2+^ efflux is sufficient to induce T3SS-dependent LTB_4_ synthesis, BMNs were pre-treated with EGTA to chelate extracellular Ca^2+^ prior to infection with *Y. pestis* T3E – if intracellular Ca^2+^ efflux is sufficient for cPLA_2_ and 5-LOX activation, then EGTA should not inhibit LTB_4_ synthesis. As observed during PLC inhibition, EGTA chelation of extracellular Ca^2+^ significantly reduced LTB_4_ production compared to untreated BMNs (Fig 4A; p≤0.0001). Influx of extracellular Ca^2+^ also requires the cell to maintain a membrane potential by efflux of intracellular potassium (K^+^) (67, 68). Therefore, if extracellular Ca^2+^ is required, disrupting the K^+^ gradient should also inhibit LTB_4_ synthesis. As predicted by EGTA treatment, increasing the extracellular K^+^ concentration significantly inhibited LTB_4_ synthesis by *Y. pestis* T3E-infected BMNs (Fig 4A; p≤0.0001). Finally, we also treated BMNs with a pharmacological inhibitor of STIM1 (SKF), which activates SOC-mediated Ca^+^ flux across the plasma membrane (69), and observed a significant decrease in LTB_4_ synthesis (Fig 4A; p≤0.0001). Together, these data support that that extracellular Ca^2+^ flux is required for the T3SS-dependent LTB_4_ synthesis, and it is mediated by the PLC-STIM1 pathway in neutrophils. Interestingly, while PLC does not appear to be required for LTB_4_ synthesis by macrophages (Fig. 3), there remains a requirement for STIM1 and extracellular Ca^2+^ flux (Fig. 4B), further indicating that alternative pathways are involved in triggering Ca^2+^ flux needed for LTB_4_ synthesis in macrophages.

**Fig. 4.**
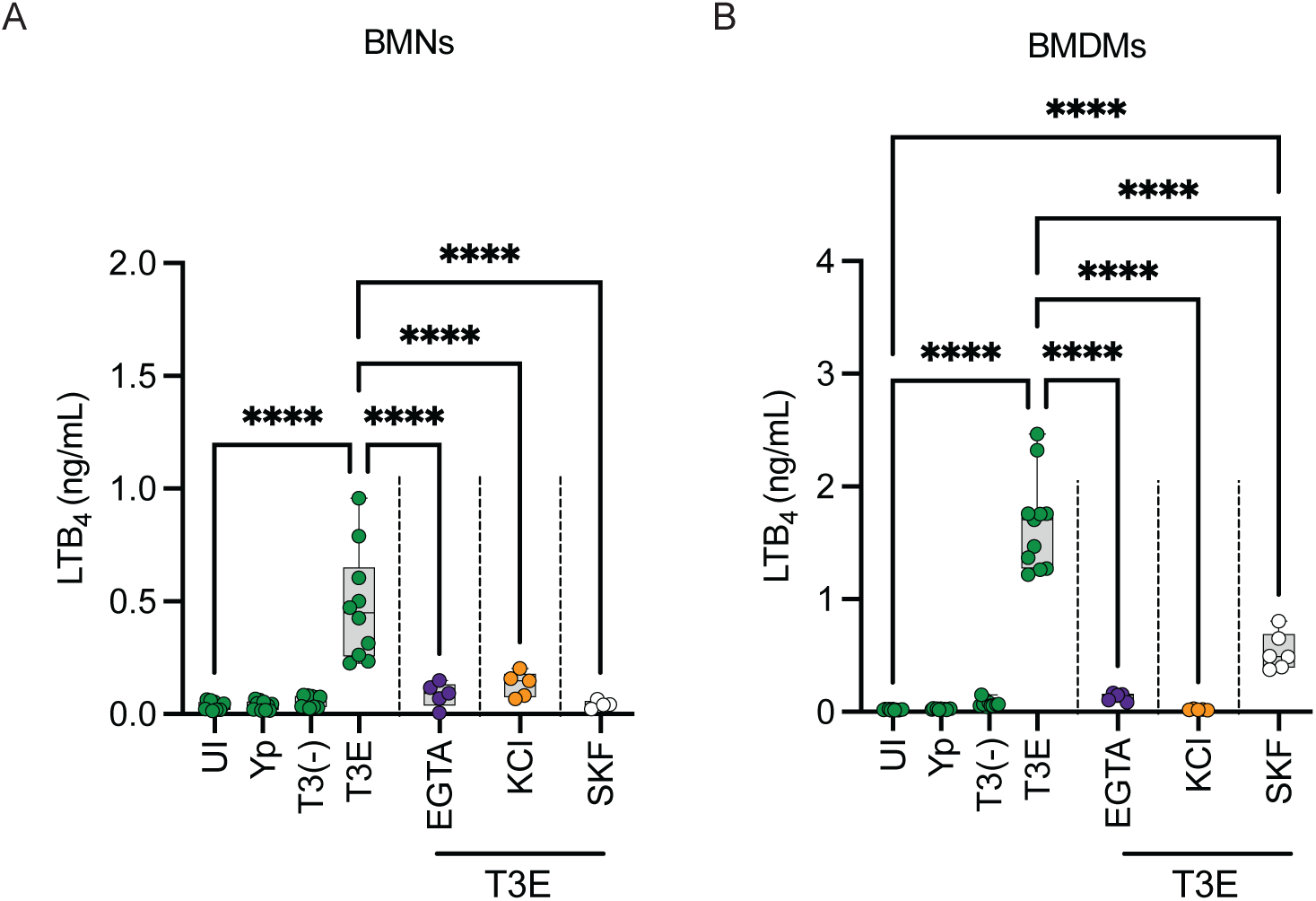
Influx of extracellular Ca^2+^ is required for *Y. pestis* T3SS-dependent LTB_4_ synthesis. (A) BMNs or (B) BMDMs were infected with *Y. pestis, Y. pestis* T3E, or *Y. pestis* T3^(−)^. Leukocytes pretreated with EGTA (1 mM; purple circles) for 30 min, with (A) 50 mM or (B) 100 mM KCl (orange circles) for 30 min, or with STIM1 inhibitor (SKF, 50 μM; white circles) for 2 min prior to infection with *Y. pestis* T3E. Leukocytes were infected at an MOI of 20 for (A) 1 h or (B) 4 h. LTB_4_ was measured from supernatants by ELISA. Each symbol represents an independent biological infection, and the box plot represents the median of the group ± the range. UI = uninfected. One-way ANOVA with Tukey’s *post hoc* test compared to each condition. ****=p≤0.0001.

### SKAP2 is required for LTB_4_ synthesis by neutrophils but not macrophages

YopH inhibits PLC-mediated Ca^2+^ flux in neutrophils during *Y. pseudotuberculosis* infection by modifying proteins of the SKAP2/SLP-76/PRAM/Vav signaling hub (28, 48, 49). Because YopH independently inhibits LTB_4_ synthesis in neutrophils, we sought to determine if this hub is also required for T3SS-dependent LTB_4_ synthesis by measuring LTB_4_ synthesis by leukocytes from SKAP2^−/−^ mice infected with *Y. pestis*, *Y. pestis* T3E, or *Y. pestis* T3^(−)^. Unlike BMNs from C57BL/6J mice, SKAP2^−/−^ BMNs did not synthesize LTB_4_ in response to any of the strains tested (Fig 5A). To ensure that SKAP2^−/−^ BMNs were not generally defective in LTB_4_ synthesis, SKAP2^−/−^ BMNs were treated with the Ca^2+^ ionophore A23187, and cells robustly produced LTB_4_ (Fig 5B). Complementing the PLC inhibitor data, SKAP2^−/−^ BMDMs were not impaired in LTB_4_ synthesis (Fig 5C; p≤0.0001). Together, these data demonstrate that T3SS-dependent LTB_4_ synthesis by neutrophils requires SKAP2 but is independent of SKAP2 in macrophages.

**Fig. 5.**
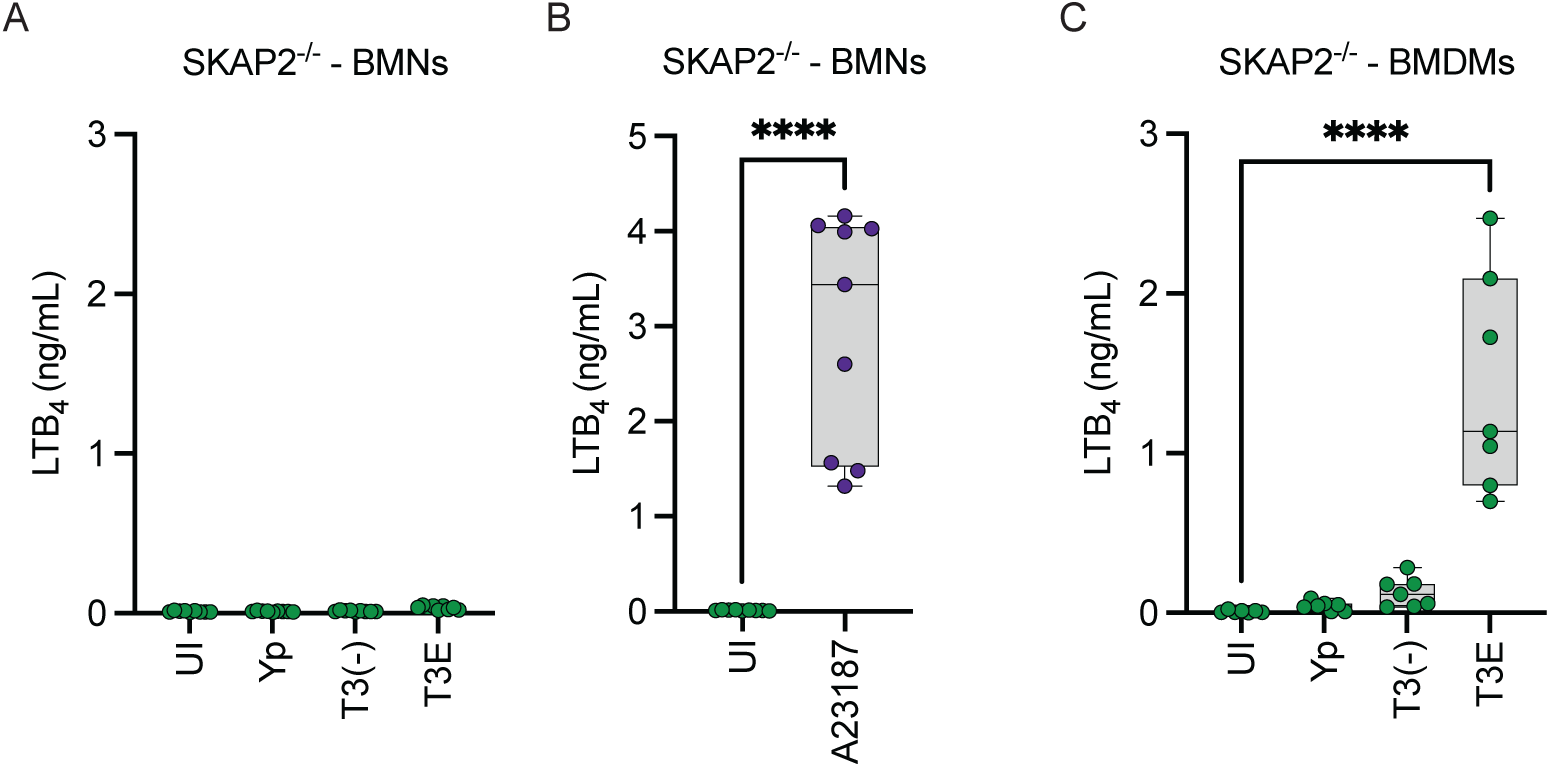
SKAP2 signaling required for LTB_4_ synthesis in BMN. SKAP2^−/−^. (A) BMNs or (C) BMDMs were infected with *Y. pestis, Y. pestis* T3E, or *Y. pestis* T3^(−)^ at an MOI of 20 for (A) 1 h or (C) 4 h. (B) SKAP2^−/−^ BMNs were treated with A23187 (1 μM; purple circles) for 1 h. (A-C) LTB_4_ was measured from supernatants by ELISA. Each symbol represents an independent biological infection, and the box plot represents the median of the group ± the range. UI = uninfected. One-way ANOVA with Dunnett’s *post hoc* test compared to uninfected. ****=p≤0.0001.

### Activation of MAPK signaling required for LTB_4_ synthesis is independent of the T3SS

In addition to Ca^2+^ flux, LTB_4_ synthesis requires MAPK signaling to phosphorylate cPLA_2_ and 5-LOX (7, 70–72). While we previously showed that p38 and ERK1/2 are phosphorylated in human neutrophils infected with a high MOI (100 bacteria/cell) of *Y. pestis* T3^(−)^ (37), the MAP kinases responsible for T3SS-dependent LTB_4_ synthesis have not been defined. Therefore, to determine whether p38 and ERK1/2 are phosphorylated during interactions with *Y. pestis* T3E, leukocytes were infected with *Y. pestis*, *Y. pestis* T3E, or *Y. pestis* T3^(−)^ at an MOI of 20. Both p38 and ERK1/2 were phosphorylated in BMNs infected with *Y. pestis* T3E but not *Y. pestis* (Fig 6A-B). For BMDMs, we observed elevated basal levels of p38 and ERK1/2 phosphorylation in uninfected cells, which has been reported in M1 polarized macrophages by others (73–76), but phosphorylation of both kinases was significantly lower in *Y. pestis* infected cells (Fig 6C-D). While p38 phosphorylation was slightly elevated in BMDMs infected with *Y. pestis* T3E, ERK1/2 phosphorylation appeared to decrease slightly, suggesting that p38 phosphorylation may be driving LTB_4_ synthesis in BMDMs. However, in both leukocytes we observed similar trends in phosphorylation in cells infected with *Y. pestis* T3E or *Y. pestis* T3^(−)^, indicating that activation of MAPK signaling is in response to a PAMP unrelated to the T3SS.

**Fig. 6.**
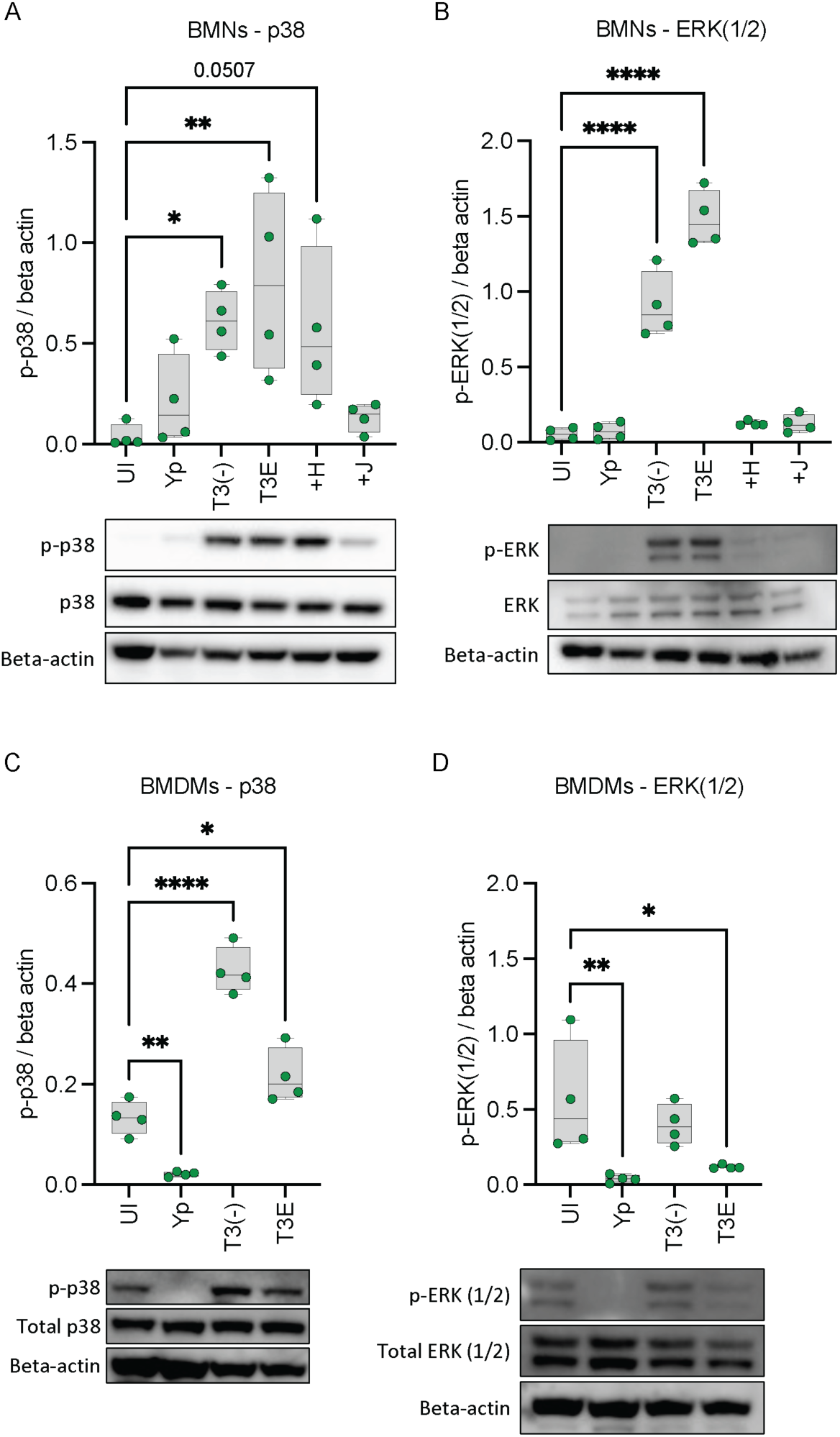
Activation of MAPK signaling required for LTB_4_ synthesis is independent of the T3SS. (A-B) BMNs or (C-D) BMDMs were infected with *Y. pestis, Y. pestis* T3E, *Y. pestis* T3^(−)^ at an MOI of 20 for (A-B) 1 h or (C-D) 30 min. (A-B) BMNs were infected *Y. pestis* TE3 expressing only one Yop effector (+H = YopH; +J = YopJ) at an MOI of 20 for 1 h. Densitometry and representative WB images for (A,C) p-p38 or (B,D) p-ERK1/2 from whole cell lysates normalized to beta actin. (A-D) Each symbol represents an independent biological infection, and the box plot represents the median of the group ± the range. UI = uninfected. One-way ANOVA with Dunnett’s *post hoc* test compared to uninfected. *=p≤0.05, **=p≤0.01, ****=p≤0.0001.

### YopH inhibits ERK phosphorylation in neutrophils

In BMNs, YopH and YopJ can independently inhibit LTB_4_ synthesis. However, both effector proteins have different targets within the host cell, which can lead to inhibiting both Ca^2+^ and/or MAPK signaling (22, 29). While we have shown that YopJ inhibition of ERK1/2 phosphorylation is sufficient to block LTB_4_ synthesis by human neutrophils (37), and Shaban et al. previously showed that YopH from *Y. pseudotuberculosis* inhibits ERK1/2 phosphorylation in neutrophils (49), whether YopH can also sufficiently inhibit ERK1/2 phosphorylation in our model has not been tested. Therefore, we measured ERK1/2 and p38 phosphorylation in BMNs infected with *Y. pestis* T3E strains expressing only YopH or YopJ. As predicted from our work with human neutrophils (37), phosphorylation of both MAP kinases was inhibited in BMNs infected with the YopJ expressing strain (Fig 6A and B; +J). However, infection with the YopH expressing strain appeared to only inhibit ERK1/2 phosphorylation and did not significantly impact p38 phosphorylation (Fig 6A and B; +H samples). These data suggest that YopH can block both Ca^2+^ and ERK1/2 phosphorylation in neutrophils during interactions with *Y. pestis*.

### Inflammasome activation enhances LTB_4_ synthesis in macrophages but not neutrophils

Previous studies with *Y. pseudotuberculosis* indicate that the T3SS translocase is recognized by NLRP3, leading to activation of the caspase 1 inflammasome and pyroptosis (39, 77). Inflammasome activation has been linked to LTB_4_ synthesis in response to some PAMPS but is dispensable for others (11, 40, 41). Therefore, to determine if the NLRP3/CASP1 inflammasome contributes to LTB_4_ synthesis in response to the T3SS, leukocytes isolated from NLRP3^−/−^ or CASP1/11^−/−^ mice were infected with *Y. pestis*, *Y. pestis* T3E, or *Y. pestis* T3^(−)^, and LTB_4_ synthesis was compared to cells isolated from WT mice. Absence of NLRP3 or CASP1/11 did not diminish the neutrophil response to *Y. pestis* T3E (Fig 7A). Furthermore, treatment of *Y. pestis* T3E infected BMNs or human PMNs with the pan-caspase inhibitor zVAD also did not decrease LTB_4_ synthesis (Fig 7B and C). In contrast, LTB_4_ synthesis was significantly lower in both NLRP3 and CASP1/11 deficient BMDMs (Fig 7D; p≤0.0001). However, LTB_4_ synthesis by *Y. pestis* T3E infected cells was still greater than uninfected, *Y. pestis*, or *Y. pestis* T3^(−)^ infected cells, indicating that the NLRP3/CASP1 inflammasome enhances the LTB_4_ response by macrophages but not neutrophils. Because CASP1/11^−/−^ macrophages still synthesized lower levels of LTB_4_ in response to the *Y. pestis* T3E strain, we next asked if this synthesis was dependent on PLC signaling, and observed only a moderate decrease in LTB_4_ when infected cells were treated with the PLC inhibitor U73122 (Fig 7E). Interestingly, we observed a much more significant decrease if we inhibited phagocytosis by treatment with cytoD or during infection with a *Y. pestis* T3E strain expressing only YopE, which is a potent inhibitor of phagocytosis (Fig 7E; p≤0.0001), suggesting that phagocytosis is key to initiating LTB_4_ synthesis in macrophages.

**Fig. 7.**
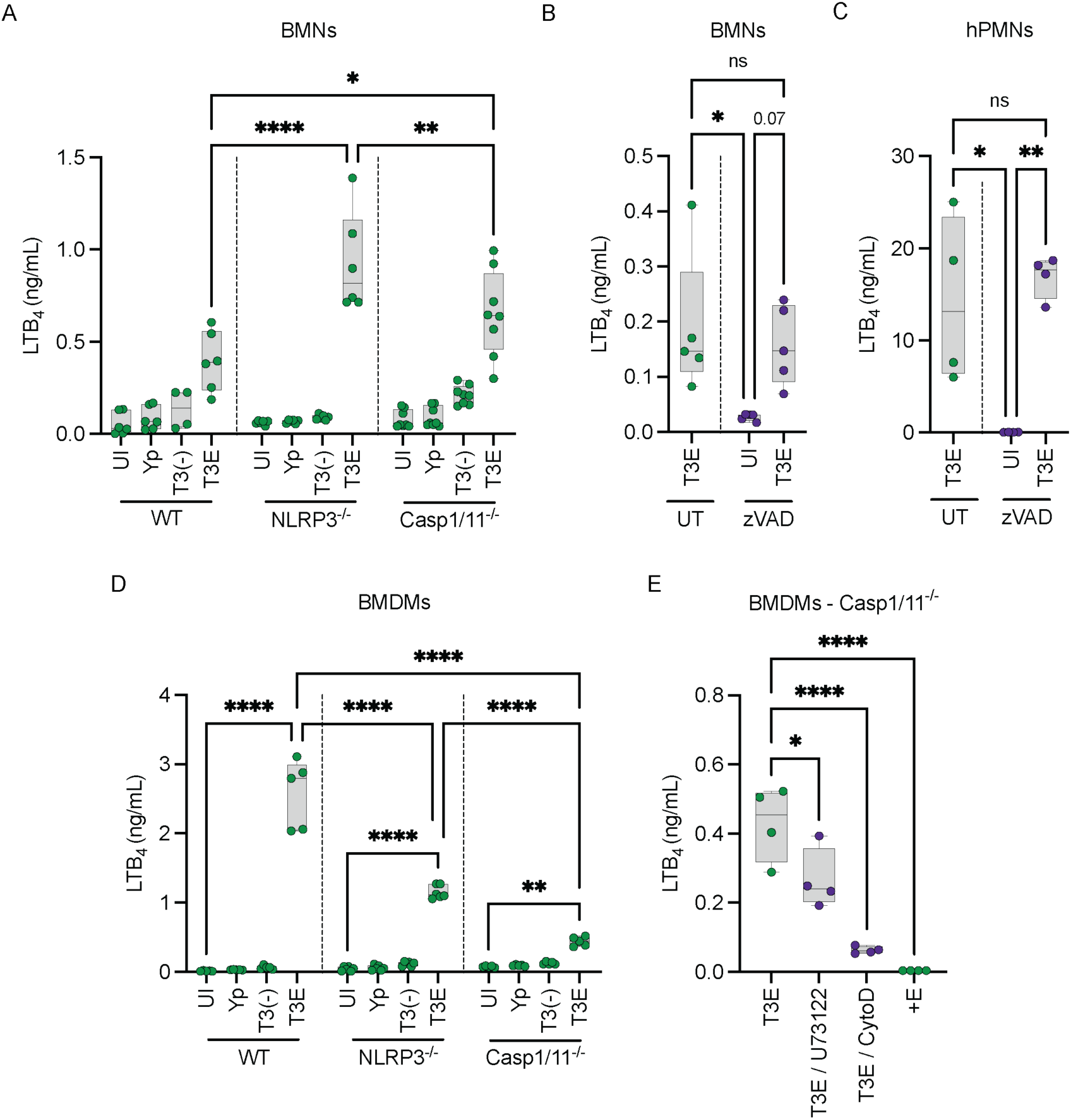
Inflammasomes enhance LTB_4_ synthesis in BMDMs but are dispensable in BMNs. (A) BMNS or (D) BMDMs from WT, NLRP3^−/−^, CASP1/11^−/−^ mice were infected with *Y. pestis, Y. pestis* T3E, or *Y. pestis* T3^(−)^. (B) C57BL/6J BMNs or (C) human PMNs (hPMNs) pretreated with zVAD inhibitor (100 μM; purple circles) for 30 min prior to infection with *Y. pestis* T3E. (E) BMDMs from CASP1/11^−/−^ mice were infected with *Y. pestis* T3E or *Y. pestis* T3E strain expressing only YopE (+E). BMDMs were either left untreated (green circles) or pretreated with PLC inhibitor (U73122, 20 μM; purple circles) or Cytochalasin D (10 μM; purple circles) for 30 min prior to infection with *Y. pestis* T3E. Leukocytes were infected at an MOI of 20 for (A-C) 1 h or (D-E) 4 h. (A-E) LTB_4_ was measured from supernatants by ELISA. Each symbol represents an independent biological infection, and the box plot represents the median of the group ± the range. UI = uninfected. UT = untreated. ns = not significant. One-way ANOVA with Tukey’s *post hoc* test compared to each condition for A-D., or with Dunnett’s *post hoc* test compared to untreated/T3E for E. *=p≤0.05, **=p≤0.01, ***=p≤0.001, ****=p≤0.0001.

## Discussion

Establishing a non-inflammatory environment during the early stages of plague is crucial for the progression of disease (20). We and others have shown that *Y. pestis* subverts the host innate immune response by inhibiting leukocyte chemotaxis (78), phagocytosis (32, 33), neutrophil degranulation (36, 37), neutrophil ROS production (33, 35), and inflammatory lipid, cytokine, and chemokine release (34, 37, 38). Despite the T3SS being a PAMP, the Yop effectors are highly efficient at preventing immune cell activation, including inhibiting the synthesis of LTB_4_ needed for a proper inflammatory host response (31, 37, 38). In this study, we sought to understand how the molecular signaling pathways stimulated by the T3SS lead to the synthesis of LTB_4_ and to define how the pathogen uses specific Yop effectors to block this response. Using these data, we have developed a working model showing a differential response to the T3SS between neutrophils and macrophages needed for LTB_4_ synthesis – with the SKAP2/PLC signaling pathway as key for a rapid response in neutrophils and the NLRP3/CASP1 inflammasome more important in macrophages (Fig 8).

**Fig. 8.**
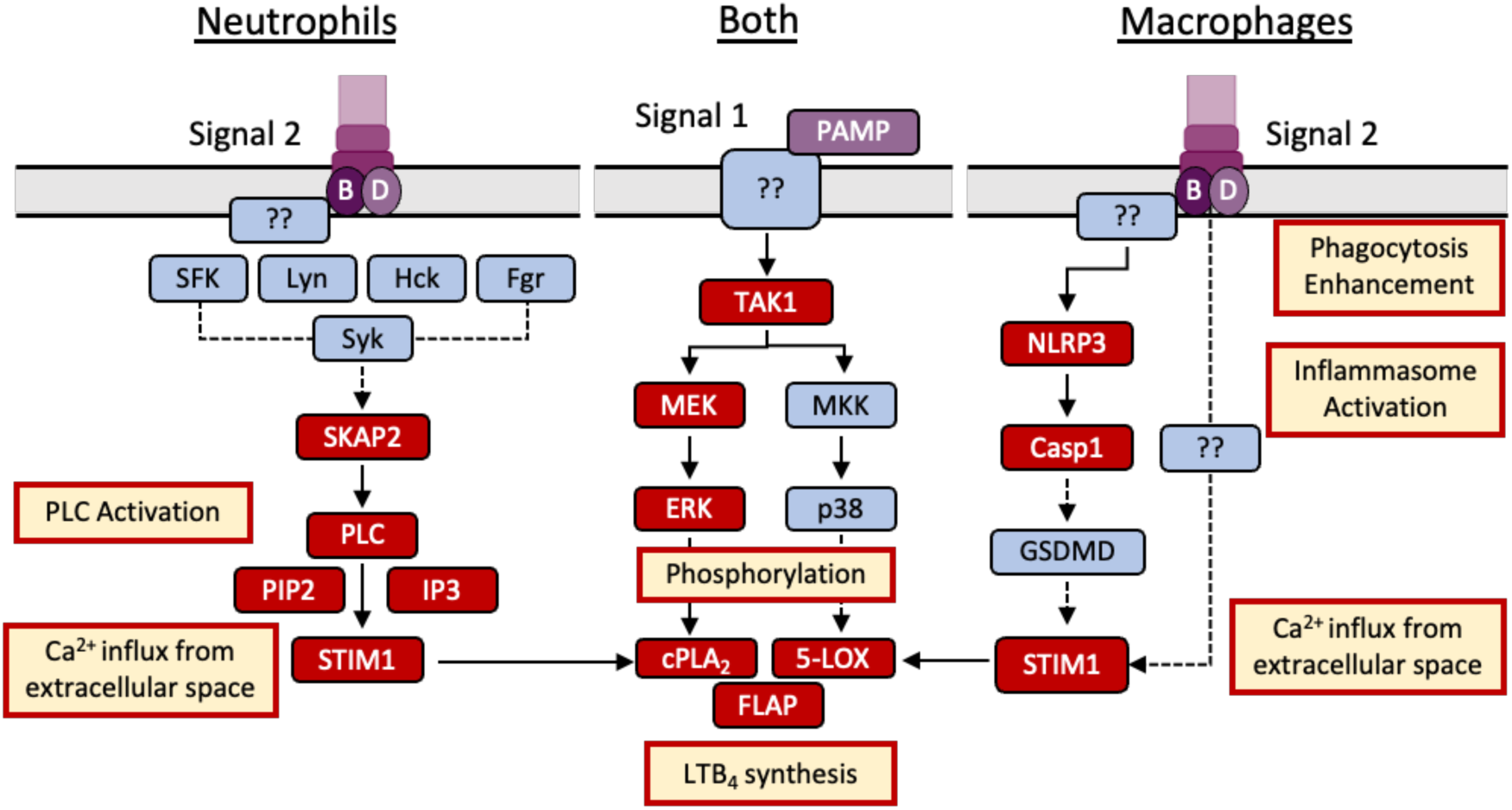
T3SS triggered LTB_4_ synthesis differs between neutrophils and macrophages. LTB_4_ synthesis requires an increase in intracellular Ca^2+^ and activation of MAPK signaling. Activation of MAPK signaling required for LTB_4_ synthesis appears to be independent of the T3SS, and instead is initiated by a currently unknown PAMP and signaling pathway(s) (Signal 1). In response to the T3SS (Signal 2), neutrophils require Ca^2+^ signaling through SKAP2/PLC/STIM1 to produce LTB_4_. Macrophages require Ca^2+^ influx from the extracellular space, with only a partial requirement of STIM1 but not SKAP2 or PLC. Inflammasome activation is not required in neutrophils for LTB_4_ synthesis but enhances the induction in macrophages. Phagocytosis also enhances LTB_4_ induction in macrophages, but not in neutrophils. Red boxes indicate factors involved in LTB_4_ synthesis examined in this study. Blue boxes indicate potential or unknown factors they may be involved but have yet to be tested.

One important discovery made here was that recognition of the T3SS leading to LTB_4_ synthesis is not specific to the *Y. pestis* T3SS. While the *S. enterica* Typhimurium SPI-1 T3SS varies structurally from the *Y. pestis* secretion system (79), *S. enterica* Typhimurium still triggers LTB_4_ in a SPI-1 T3SS-dependent manner (Fig 1). Additionally, LTB_4_ synthesis by *S. enterica* Typhimurium infected leukocytes appears to be more responsive to the SPI-1 T3SS than the SPI-2 system. Like the *Y. pestis* T3SS, the SPI-1 T3SS engages with the host cell through the plasma membrane, while SPI-2 engages through the *Salmonella* containing vacuole (80, 81), suggesting that the host cells have evolved to sense T3SS interactions across the plasma membrane and respond by synthesizing LTB_4_. This scenario is attractive, as interactions with the plasma membrane represent the earliest interaction that leukocytes would have with pathogens and allow for rapid synthesis of the lipid as soon as contact is made. It also appears that unlike *Y. pestis*, *S. enterica* Typhimurium has not evolved effector proteins to inhibit this response, or at minimum, not to the degree that the *Y. pestis* Yop effectors can inhibit LTB_4_ synthesis, as the WT *S. enterica* Typhimurium LT strain produces LTB_4_ at levels similar to the *Y. pestis* T3E mutant. This difference supports that the inhibition of LTB_4,_ and initiation of the inflammatory cascade is an important aspect in the virulence and lifestyle of *Y. pestis* that is not required for *S. enterica* Typhimurium.

A key virulence strategy mediated by the T3SSs of both *Y. pestis* and *S. enterica* Typhimurium is the manipulation of phagocytosis (46, 82). Because phagocytosis of crystalline silica is required for LTB_4_ synthesis (11), determining the contribution of phagocytosis to LTB_4_ synthesis was another critical aspect of understanding the leukocyte response to these pathogens. In neutrophils, phagocytosis was not required for LTB_4_ synthesis, and cytochalasin D treatment induced even greater LTB_4_ synthesis in response to *S. enterica* Typhimurium. However, inhibiting phagocytosis reduced LTB_4_ synthesis by macrophages, providing the first evidence that the host cell molecular mechanisms leading to cPLA_2_ and 5-LOX activation and LTB_4_ synthesis differ between these two cell types. It is important to note that for both bacteria, LTB_4_ synthesis by macrophages was not completely inhibited by cytochalasin D treatment, indicating that synthesis is not completely dependent on phagocytosis in macrophages. Moreover, data from infections with the *Y. pestis* T3^(−)^ mutant, which is as readily phagocytosed as the *Y. pestis* T3E mutant, indicates that phagocytosis alone is not sufficient to trigger LTB_4_ synthesis in the absence of the T3SS.

Ca^2+^ flux is a critical step in the enzyme activation required for LTB_4_ synthesis (7, 14), and thus understanding the mechanisms leading to Ca^2+^ flux during host cell interactions with the T3SS expressing bacteria is key to understanding how cells are sensing this PAMP. YopB and YopD insert into the plasma membrane to form a pore needed for effector translocation into the host cell (51–53). Others have shown that the pore formed by the translocase can result in the diffusion of molecules larger than Ca^2+^, but YopN appears to limit diffusion through the YopB/D translocase (83). While the *Y. pestis* T3E strain used in these studies retains YopN, it is still possible that Ca^2+^ diffusion through the translocase could occur in infected cells. However, using pharmacological inhibitors, we have shown that PLC and store-operated Ca^2+^ signaling, and not diffusion of Ca^2+^ through the translocase pore, is likely the primary driver of Ca^2+^ flux needed for LTB_4_ synthesis in neutrophils. This is further supported by the ability of YopH to inhibit LTB_4_ synthesis, which blocks Ca^2+^ flux by inhibiting PLC phosphorylation and signaling and not ion diffusion through the translocase (28, 49), and the requirement for SKAP2, which activates PLC in response to signaling from a variety of tyrosine kinase receptors (84, 85). Identifying which tyrosine kinase receptors are phosphorylating SKAP2 in response to the T3SS is an ongoing research direction for us and will help us to better understand the mechanisms directly sensing the T3SS and/or YopB/D translocase. Contrarily, because PLC pharmacological inhibition and YopH had minimal effects on LTB_4_ synthesis in macrophages, the contribution of Ca^2+^ diffusion through the translocase in macrophages is less clear. It appears that LTB_4_ synthesis in macrophages still requires activation of STIM1 and store-operated calcium channels (Fig. 5), indicating that diffusion of Ca^2+^ through the pore is not sufficient to activate cPLA_2_ and 5-LOX in macrophages, but also that the signal to activate STIM1 is independent of PLC. PLC-independent activation STIM1 has been reported, but activation requires Ca^2+^ store depletion from the ER (86, 87), suggesting that any Ca^2+^ diffusion via the translocase pore is not directly responsible for STIM1 activation. Currently it is unclear how STIM1 is activated during macrophage interactions with the *Y. pestis* T3E strain and additional research is needed to completely understand the PLC-independent pathway(s) activated in macrophages.

In addition to Ca^2+^ flux, cPLA_2_ and 5-LOX phosphorylation via MAPK signaling is also essential for LTB_4_ synthesis (7, 70–72). Depending on the stimulus, 5-LOX phosphorylation is mediated by p38, ERK1/2, or JNK (11, 70, 88, 89), but while p38 and ERK1/2 are both phosphorylated in human neutrophils during interactions with *Y. pestis* T3E, only ERK1/2 was essential for LTB_4_ synthesis (37). While we confirmed that both p38 and ERK1/2 are also phosphorylated in murine neutrophils and macrophages (Fig 6), we also discovered that MAPK activation is not dependent on the presence of the T3SS, as we observed significant phosphorylation of the kinases in cells infected with the *Y. pestis* T3^(−)^ strain that lacks the pCD1 plasmid and the entire T3SS. These data indicate that a different PAMP(s) is triggering the MAPK pathway and that the host factors recognizing the T3SS and triggering Ca^2+^ flux differ from those required for MAPK phosphorylation. A requirement for two signals is an important check point for other responses to bacteria, for example inflammasome activation, to limit premature inflammatory responses that could be detrimental to the host if stimulated too easily. Thus, to avoid chronic inflammation, a similar two signal check point may have also evolved to limit premature LTB_4_ responses to bacteria. It is also currently unclear if this is specific for bacteria with T3SSs. Studies with other pathogens are needed to determine if multiple PAMPs/signals are also required for LTB_4_ responses to bacteria that do not express T3SSs. Importantly, these data support that the primary signal governing whether LTB_4_ is produced in recognition of the T3SS is Ca^2+^ flux, not the MAPK activation.

While BLT1-LTB_4_ signaling has been shown to enhance inflammasome activation in gout (90), asthma (6), and *Staphylococcus aureus* skin infection models (91), evidence in the literature that inflammasome activation induces LTB_4_ synthesis is limited. However, Von Moltke et al. showed that direct delivery of the NAIP5 ligand flagellin to the cytosol of resident peritoneal macrophages, but not M2 polarized BMDMS, triggers leukotriene and prostaglandin synthesis, which was dependent on NLRC4, NAIP5, and CASP1 (40). Moreover, they demonstrated that resident peritoneal macrophages produce prostaglandin E2 (PGE_2_) in response to *S. enteric* Typhimurium, but they did not report LTB_4_ synthesis or test dependence on the SPI1 T3SS (40). The new data reported here further support that polarization or activation state significantly impacts the potential of macrophages to synthesize eicosanoids during infection. Moreover, we have now identified a second example of a PAMP that triggers eicosanoid synthesis in an inflammasome-dependent manner and supports that inflammasome activation not only results in the release of protein mediators of inflammation (i.e., IL-1β and IL-18) but also lipid mediators of inflammation. The mechanisms leading to T3SS-dependent inflammasome-mediated LTB_4_ synthesis are still unclear, but the MAPK data suggests that it is involved in triggering Ca^2+^ flux, either directly or through the activation of STIM1 and the store-operated calcium channels.

In conclusion, we have shown that leukocytes have evolved to recognize the T3SS as part of two signal cascade that leads to LTB_4_ synthesis during bacterial interactions. However, the molecular mechanisms resulting in LTB_4_ synthesis differ between neutrophils and macrophages, the former dependent on the SKAP2/PLC signaling pathway and the latter on the inflammasome. Together, these data provide us with a better understanding of the early response of leukocytes to bacterial pathogens and provide an example of how a pathogen, in this case *Y. pestis*, has evolved mechanisms to subvert these responses and evade the immune recognition to cause disease.

## Material and Methods

### Ethics statement

All animal work was approved by the University of Louisville Institutional Animal Care and Use Committee (IACUC Protocol #22157). Use of human neutrophils was approved by the University of Louisville Institutional Review Board guidelines (IRB #96.0191) and written consents for use were obtained.

### Bacterial strains

Bacterial strains used in this study are listed in S1 Table. *Y. pestis* was cultured with BHI broth for 15-18 h at 26°C in aeration. Cultures were then diluted 1:10 in fresh, warmed BHI broth containing 20 mM MgCl_2_ and 20 mM Na-oxalate and cultured at 37°C for 3 h with aeration to induce expression of the T3SS. *S. enterica* Typhimurium was cultured with LB broth for 15-18 h at 37°C in aeration. Cultures were then diluted 1:10 in fresh, warmed LB broth and cultured at 37°C for 3 h with aeration to reach logarithmic growth. Bacterial concentrations were determined using a spectrophotometer and diluted to desired concentrations in fresh medium.

### Cell isolation and cultivation

Murine leukocytes were isolated from bone marrow of 7-12-week-old mice that were either C57BL/6J, C57BL/6J Tyr^−/−^ (B6 albino; Jackson Laboratory 000058), C57BL/6J Tyr^−/−^ NLRP3^−/−^, C57BL/6J Tyr^−/−^ Caspase1/11^−/−^, or BALB/c SKAP2^−/−^. Murine neutrophils were isolated using an Anti-Ly-6G Microbeads kit (Miltenyi Biotec Cat. No. 130-120-337) per the manufacturer’s instructions. Neutrophil isolations yielded ≥ 95% purity and were used within 1 h of isolation. Macrophages were differentiated from murine bone marrow in DMEM supplemented with 1 mM Na-pyruvate, and 10% FBS for 6 days. Macrophages were polarized with 20 ng/mL of GM-CSF (Kingfisher Biotech Cat. No. RP0407M) throughout the differentiation. The medium was replaced on days 1 and 3 (adapted from (92)). Human neutrophils were isolated from the peripheral blood of healthy, medication-free donors, as described previously (93). Neutrophil isolations yielded ≥ 95% purity and were used within 1 h of isolation.

### Leukocyte infections

Human neutrophils were cultured in Kreb’s buffer (w/ Ca^2+^ & Mg), mouse neutrophils (BMNs) were cultured in RPMI + 5% FBS, and macrophages (BMDMs) were cultured in DMEM + 10% FBS. Human neutrophils were adhered to 24-well plates for 30 min that were coated with pooled human serum prior to infection. BMNs were adhered to 24-well plates for 30 min that were coated with FBS prior to infection. Plates were washed twice with 1 x DPBS prior to plating the cells. BMDMs were adhered to 24-well plates 1 day prior to infection. Leukocytes were infected at a multiplicity of infection (MOI) of 20 and incubated for 1 h (neutrophils) or 4 h (macrophages) at 37°C with 5% CO_2_. All infections were synchronized by centrifugation (200 x g for 5 min). Supernatants and cells were then collected from wells to an Eppendorf tube and centrifuged for 1 min at 6,000 x g. Supernatants devoid of cells were transferred to a fresh tube and stored at −80°C until ELISA. When applicable, cell pellets were directly prepped for western blot analysis.

### Treatments and inhibitors

Prior to infection, leukocytes were treated with the following for the times and concentrations indicated in the figure legends: phagocytosis inhibitor cytochalasin D (VWR; Cat. No. 100507-376), calcium ionophore A23187 (Sigma-Aldrich; Cat. No. C7522), PLC inhibitor U73122 (Abcam; Cat. No. ab120998), STIM1 inhibitor SKF-96365 (VWR; Cat. No. 89156-792), 1 mM EGTA, 50 mM or 100 mM KCl, or the pan-caspase inhibitor Z-Vad-FMK (Enzo; Cat. No. ALX-260-020). At the time of infection, bacteria were added for a 500 μL final volume.

### Measurement of LTB_4_ by enzyme-linked immunosorbent assay

Supernatants of neutrophils and macrophages were collected and measured for LTB_4_ by ELISA per manufacturer’s instructions (Cayman Chemicals; Cat. No. 502390).

### Western blots

Pellets were lysed over ice in 1x Novex lysis buffer and processed through Qiashredders (Qiagen; Cat. No. 79654). Samples were boiled for 10 min, and 10 μL was separated on a 10% SDS-PAGE gel. Samples were immunoblotted with polyclonal anti-p-p38 (Cell Signaling; Cat. No. 9211S), anti-p38 (Cell Signaling; Cat. No. 9228), anti-p-p44/42 (ERK1/2) (Cell Signaling; Cat. No. 9101s), anti-p44/42 (Cell Signaling; Cat. No. 4696), anti-beta-actin (Cell Signaling; Cat. No. 3700s). All antibodies were diluted 1:1000 or except anti-p44/42, which was diluted 1:2,000. Anti-rabbit (Sigma-Aldrich; Cat. No. A9169) or anti-mouse (ThermoFisher Scientific; Cat. No. 31430) IgG HRP secondary antibodies were diluted to 1:20,000. SuperSignal West Femto maximum-sensitivity substrate (ThermoFisher Scientific; cat. no. 34095) was used to detect antigen-antibody binding. Densitometry was performed using ImageJ software to quantify bands, normalized to beta actin.

### Confocal microscopy

BMNs infected at an MOI of 10 with a GFP expressing *Y. pestis* T3E strain were pretreated with 10 μM cytochalasin D or DPBS. After 1 h of infection, cells were then fixed with 4% PFA (Sigma Aldrich; cat. no. P6148-500G), blocked with 3% BSA-PBS (Sigma; A4503-100G), stained with rabbit anti-*Yersinia pestis* sera (1:1,000; lot UL25, 9/14/2013) overnight at 4°C, followed by donkey anti-rabbit Alexa Fluor 647 (1:1,000; JacksonImmuno Research; cat. no. 711-605-152) for 2 hours at room temperature, and finally with Hoechst (1:350; cat. no. ThermoScientific; 62249) at room temperature for 15 minutes. Cells were then mounted in Prolong Gold (Invitrogen; cat. no. P36980) and visualized with z-stack images using a confocal Olympus Fluoview FV3000 UPlanxApo. To quantify the rates at which bacteria were phagocytosed, 3D volume Pearson correlation coefficients were calculated for eGFP and Alexa647.

### Statistics

For all studies, male and female mice or human donors were used and no sex biases were observed for any phenotype. For all experiments, each data point represents data from biologically independent experiments performed on different days. Where appropriate and as indicated in the figure legends, statistical comparisons were performed with Prism (GraphPad) using one-way analysis of variance (ANOVA) with Dunnett’s or Tukey’s *post hoc* test, or T-test with Mann-Whitney’s *post hoc* test. P values ≤ 0.05 were considered statistically significant and reported.

## Acknowledgements

The authors would like to thank Dr. Jon Goguen for sharing *Y. pestis* strains and Dr. Micah Worley for *S. enterica* Typhimurium strains used in these studies. We would also like to acknowledge Pathricia “Angel” Leus and Dr. Joan Mecsas for generously providing us with bone marrow from SKAP2^−/−^ mice and Dr. Jonathan Warawa for sharing Tyr^−/−^, NLRP3^−/−^, and CASP1/11^−/−^ mice.

## Supplementary Info

**Figure S1.**
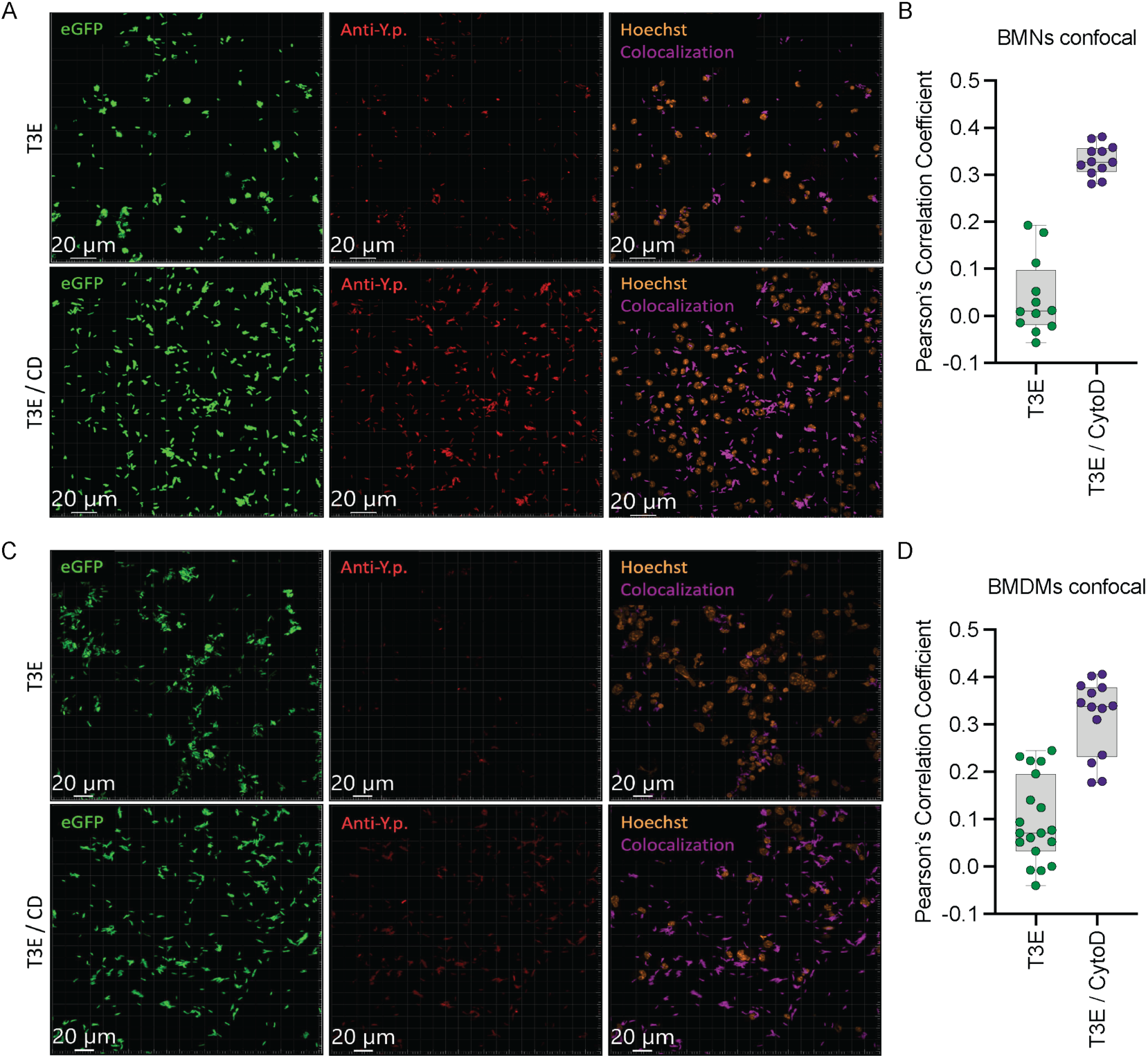
Cytochalasin D successfully inhibits BMN and BMDM phagocytosis of *Y. pestis*. (A-B) BMNs or (C-D) BMDMs were either left untreated (T3E) or pre-treated with cytochalasin D (10 μM; T3E / CD) for 30 min prior to infection with *Y. pestis* T3E at an MOI of 10 for (A-B) 1 h or (C-D) 4 h. (A,C) Representative confocal images of 3 biological replicates; eGFP=intracellular bacteria; Anti-Y.p.=extracellular bacteria. (B,D) Pearson scores calculated for eGFP and Anti-Y.p.-Alexa647 from 3 biological replicates, 3-4 images collected from each replicate.

**Table S1.**
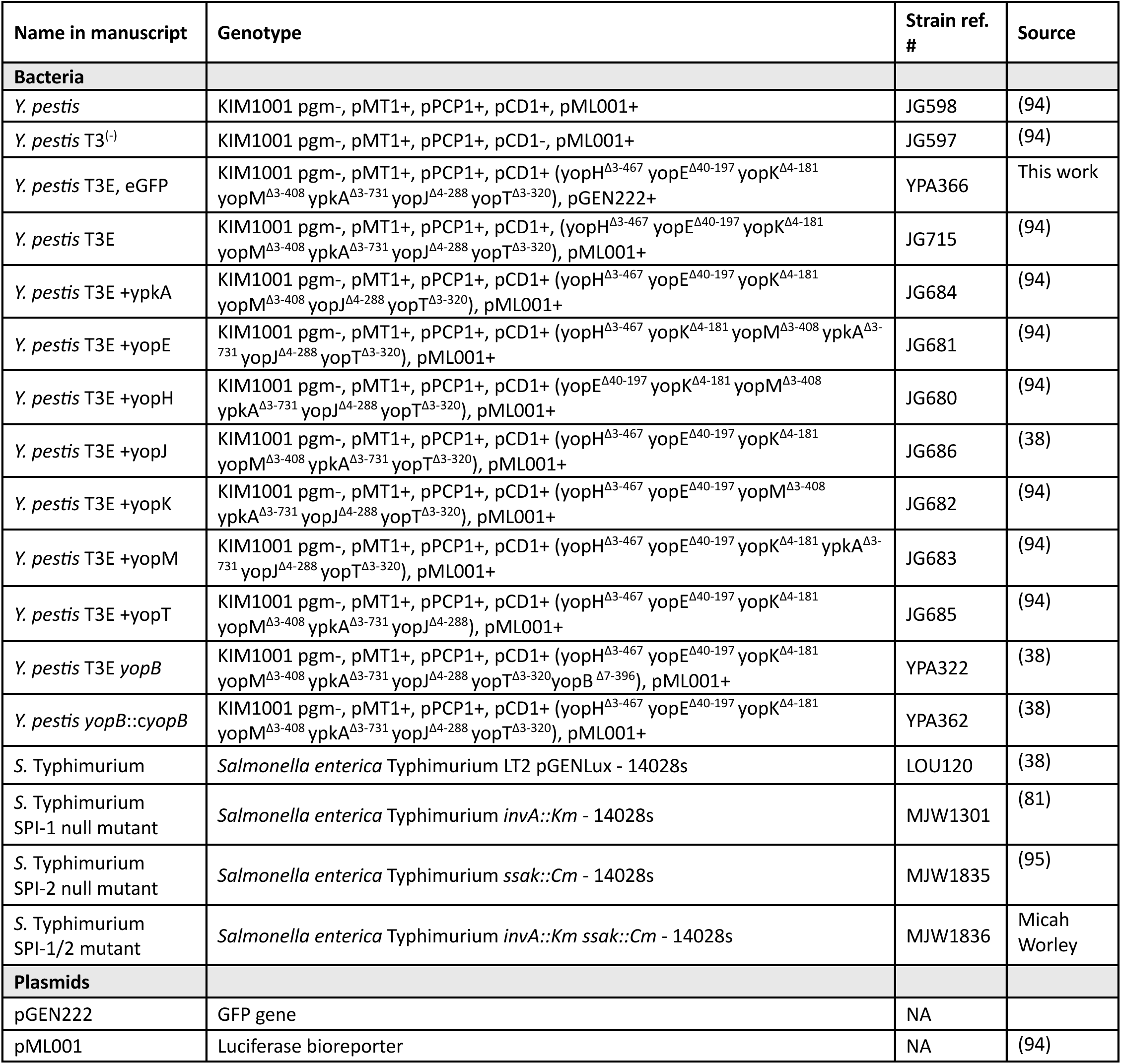
Bacterial strains and plasmids used in this study.

